# Amygdala Self-Neuromodulation Capacity as a Window for Process-Related Network Recruitment

**DOI:** 10.1101/2024.06.06.592364

**Authors:** Guy Gurevitch, Nitzan Lubianiker, Taly Markovits, Ayelet Or-Borichev, Naomi B. Fine, Tom Fruchtman-Steinbok, Jacob N. Keynan, Alon Friedman, Neomi Singer, Talma Hendler

## Abstract

Neurofeedback (NF) has emerged as a promising avenue for demonstrating process-related neuroplasticity, enabling self-regulation of brain function. NF targeting the amygdala has drawn attention for therapeutic potential in psychiatry, by potentially harnessing emotion-regulation processes. However, not all individuals respond equally to NF training, possibly due to varying self-regulation abilities. This underscores the importance of understanding the mechanisms behind successful neuromodulation (i.e. capacity). This study aimed to investigate the establishment and neural correlates of neuromodulation capacity by using data from repeated sessions of Amygdala Electrical Finger Print (EFP)-NF and post-training fMRI-NF session.

Results from 97 psychiatric patients and healthy participants revealed increased amygdala-EFP neuromodulation capacity over training, associated with post-training amygdala fMRI modulation-capacity and improvements in alexithymia. Individual differences in this capacity were associated with pre-training amygdala reactivity and initial neuromodulation success. Additionally, amygdala down-regulation during fMRI-NF co-modulated with other regions such as the posterior-insula and parahippocampal gyrus. This combined modulation better explained EFP-modulation capacity and improvement in alexithymia than the amygdala modulation alone, suggesting the relevance of this broader network to the gained capacity. These findings support a network-based approach for NF and highlight the need to consider individual differences in brain function and modulation capacity to optimize NF interventions.

## Introduction

One of the strongest, albeit challenging ways to demonstrate process-related neuroplasticity is to manipulate an assumed neural mechanism of this process, identify associated neural modification, and test their effect on behavioral and/or psychological indication of this process. Neurofeedback (NF), a non-invasive brain-computer-interface technique, has gained increasing attention in recent years as a potential tool for demonstrating process related neuroplasticity through self-neuromodulation of one’s brain activity and connectivity. It was further shown that with repeated sessions of training, this reinforcement-based procedure results in an increasing **capacity** for self-neuromodulation (i.e. defined herby as the difference in activation between ’Rest’ and ’Regulate’ conditions, see Figure 1)[1,2].

**Figure 1.**
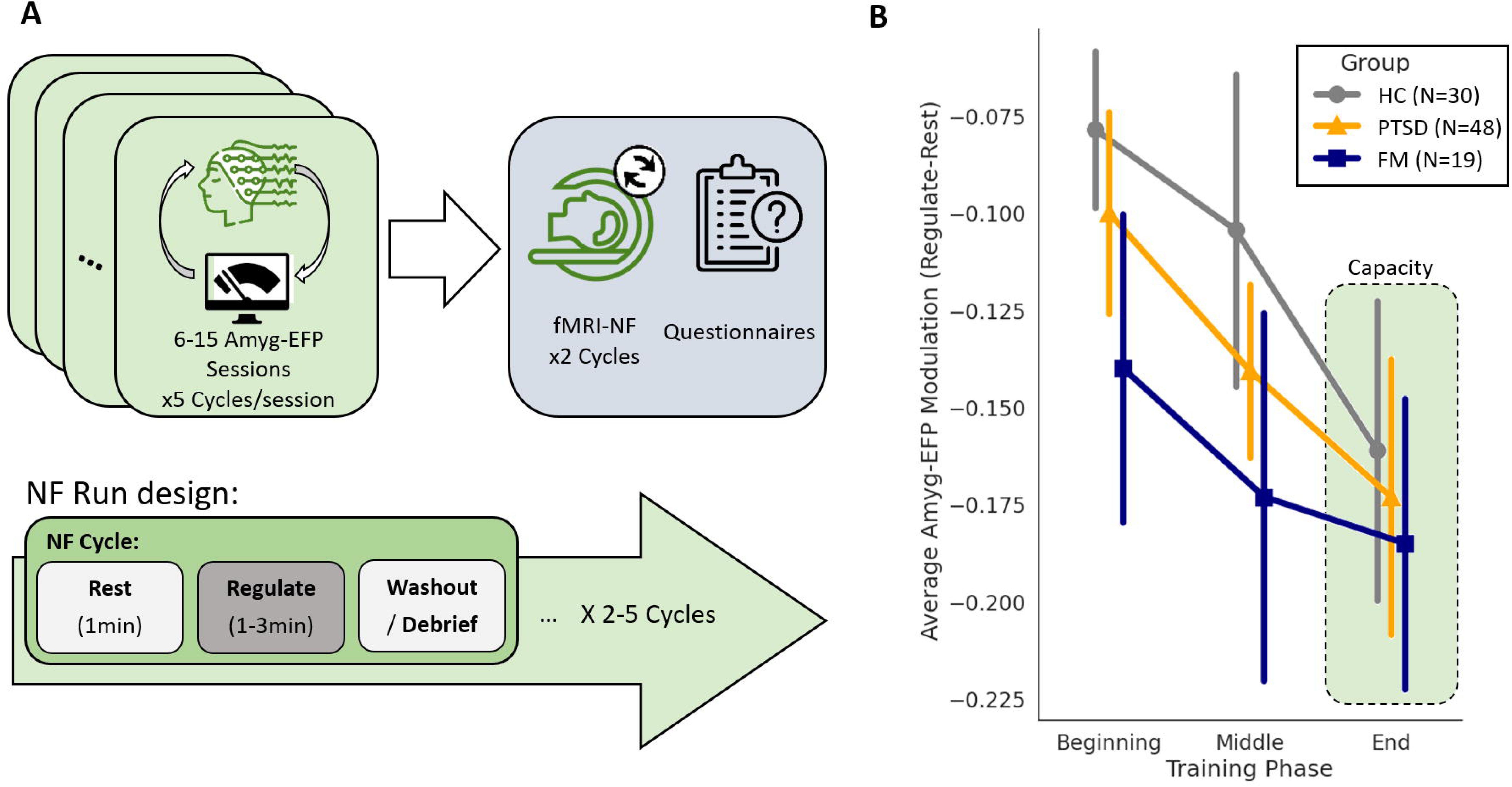
Study design and Amyg-EFP Capacity overview. (A) A schematic illustration of the study design. Participants from three subgroups of healthy control, PTSD patients and Fibromyalgia patients went through Amyg-EFP-NF training protocols, followed by post-training assessment which included fMRI-NF and questionnaires. Each Amyg-EFP session included 5 cycles combined of interleaved ’Rest’ and ’Regulate’ conditions. (B) Average Amyg-EFP modulation across the training phases, showing a significant main effect for training. Group differences were not significant. The HC group is shown in gray circles, PTSD in orange triangles and FM in blue squares. The green highlight shows the final Amyg-EFP capacity used in further analyses.

A leading target for neuromodulation in psychiatry has been the Amygdala; a major limbic node widely connected to cortical and subcortical regions, which is accordingly involved in multiple mental processes such as fear response[3], anxiety[4,5] and emotion regulation[6,7]. Correspondingly, abnormalities in amygdala activity and its connectivity with other brain areas have been suggested as trans-diagnostic markers in psychiatric disorders[8] that are observed, for example, in major depressive disorder[9,10], anxiety disorders[11,12], borderline personality disorder (BPD)[13] and post-traumatic stress disorder (PTSD)[14,15].

The feasibility and utility of Amygdala NF have recently been examined in a meta analytic review, which showed that individuals can learn to voluntarily change their amygdala activity in real-time, and that multiple sessions of Amygdala NF have resulted in improved clinical status among patients with depression, anxiety and PTSD [16]. Importantly, it was further reported that Amygdala כMRI-NF led to neural modifications relevant to the emotion regulation process, such as reduced amygdala reactivity to emotional stimuli[17–22] and enhanced functional connectivity between the amygdala and medial prefrontal cortex (mPFC), premotor cortex, and rostral anterior cingulate cortex[23–25].

However, it is also well recognized that not all participants are able to regulate their brain activity, including the amygdala, to a similar extent, and that the degree of success in NF training might predict behavioral or clinical change[26–29]. Indeed, meta-analytic reports of the NF literature reveal significant individual differences in neuromodulation capacity (also referred to as regulation success), with percentages of poor-capacity ranging from 30-50% in fMRI-NF studies[16,30] and up to 57% in EEG-NF studies[31].

Understanding the mechanisms of successful self-neuromodulation not only contributes basic scientific understanding of NF mechanisms but could also inform tailored training strategies that may optimize individual neuromodulation capacity and expand the range of effective clinical applications for diverse patient groups within a disorder category[32]. This could further improve the utilization of Amygdala NF [33,34].

Previous accounts of the psychological factors that might affect modulation success have highlighted sustained attention and motivation as crucial factors for desired modulation[33], while difficulties in identifying and regulating emotions – as indicated by alexithymia, may hinder it[35]. Methodological factors, such as the type and nature of the interface[36] or the type of instructions[37], were also found to influence modulation success. However, these may depend on the neural target and population studied. While some accounts of neural predictors of neuromodulation success have been made, specifically regarding the target activation levels before training [34], the precise neurobehavioral mechanisms underlying the aspect of the capacity of self-neuromodulation through NF is yet to be unraveled. Accordingly, it has been debated whether a-priori neuropsychological conditions (e.g. level of activation, ability to concentrate) or self-generated acquired skill determine the neuromodulation capacity[30,31,38].

Research has shown that successful neuromodulation was also accompanied by structural[39] and functional[40] neural alterations. fMRI studies have further identified specific brain regions and networks that are involved in self-neuromodulation extending beyond the target region, such as the prefrontal cortex, anterior cingulate cortex, and insula[16,41]. These regions have been implicated as the underlying “general neural processes” in NF across targets. However, it is yet unclear if these regions also determine the range of neuromodulation capacity over training. We recently showed that successful vs. unsuccessful down regulation of amygdala activity during one fMRI-NF session involved a restricted set of regions extending beyond the neural target, and not related to the aforementioned general NF processes, such as the posterior insula, precuneus and ventromedial PFC[16]. However, these findings are based on a single NF session, which is limited in its ability to uncover the full potential of modulation capacity that may be revealed with repeated training.

The most precise way of modulating the amygdala is via fMRI-NF. However, fMRI is costly and largely inaccessible outside of hospital settings, which makes it not scalable. This limitation hinders the optimization of training in terms of the number of sessions and the context in which training takes place. Another established way for targeting neural activity with high temporal precision is EEG, an accessible and affordable neurophysiological imaging technology. However, the neuroanatomical precision of EEG is questionable, resulting in poor target localization, particularly in deep brain region as the amygdala. To advance beyond the state of the art, we have established an analytic way to benefit from both technologies, resulting in fMRI informed EEG models[42–44]. Such a one-class prediction model is based on simultaneous acquisition of fMRI and EEG in an independent dataset, and the prediction of well localized fMRI activity in the amygdala from the EEG signal; termed herby Amygdala Electrical Fingerprint (Amyg-EFP). Using such a scalable neural target we were able to demonstrate that after 10-15 training sessions with Amyg-EFP-NF, PTSD patients significantly improved in their clinical status[45–47], Fibromyalgia patients showed improvement in symptom burden[29], and after six sessions, healthy controls under chronic stress showed improved alexithymia scores[23]. However, none of these studies was designed or powered enough to examine the underlying neural mechanisms of successful EFP modulation capacity.

Building on this accumulated data of patients and healthy participants undergoing both repeated sessions of Amyg-EFP-NF (for down regulation) and an outcome session of Amygdala fMRI-NF, the current study pursued two aims. The first aim was to characterize the establishment of *neuromodulation capacity* and to assess its process-relevance following training for amygdala down-regulation, as well as its influence by individual differences in neural reactivity and flexibility. We hypothesized that EFP modulation capacity will enhance over training and that this capacity will be associated with neural and mental outcomes; specifically, fMRI-NF modulation and alexithymia, respectively. We further expected that neural activity or modulation success at the beginning of training will contribute to the established neuromodulation capacity at the end of training. The second aim was to reveal the neural correlates of successful neuromodulation (i.e. high capacity) by assessing the involvement of regions beyond the target area of modulation during fMRI-NF. We hypothesized that high capacity of amygdala down-modulation via fMRI-NF will be accompanied by similar modulation in additional regions as previously shown on healthy participants[16], and that by adding these regions to the target modulation, the variance in EFP capacity and its related process change of alexithymia will be better explained than by the amygdala modulation alone.

## Materials and Methods

### Participants

In this project, we reanalyzed raw data from several studies that took place in the Sagol Brain Institute at Tel-Aviv Sourasky Medical Center between the years 2015-2021. All participants gave written informed consent and the studies were conducted under the approval of the institutional review board. The healthy controls (HC) group included thirty male Israeli Defense Forces (IDF) combat soldiers undergoing basic military training who were a subgroup from a study by Keynan et al., 2019[23]. These participants were a part of the Amyg-EFP-NF group in that study that also underwent post-training fMRI. The PTSD group included forty-eight patients who were clinically interviewed and met the Clinician Administered PTSD Scale (CAPS-5)[48] criteria for post-traumatic stress disorder (PTSD). Eighteen patients from a study by Fructhman at al., 2021[46], twenty patients from a study by Fine et al., 2023[45], and ten patients from an unpublished cohort who met the screening criteria but underwent a shorter training protocol (six sessions). The FM group included nineteen patients with a confirmed diagnosis for fibromyalgia (FM) according to the American College of Rheumatology (ACR) 2010 criteria[49] from a study by Or-Borichev et al., (in preparation). Other participants from the aforementioned studies were excluded if they were in the study’s control groups (i.e. No-NF) or if they failed to complete at least 2/3 of the assigned training regimen including the post-training fMRI session. Group demographics and personality scores at baseline are described in supplementary table 1.

### Amyg-EFP Training

After enrolling and signing informed consent, participants were assigned an Amyg-EFP-NF training regimen according to the specific study protocol. Each NF session lasted 40-60 minutes including preparation time and began with a three-minute baseline recording at rest. Next, each participant performed five NF cycles, comprising three consecutive conditions – passively attending the interface (’Rest’, 1 min), downregulating Amyg-EFP (’Regulate’, 1-3 min) and debriefing by the experimenter which included a performance summary graph and verbal description of the mental strategies employed during that cycle (Figure 1A). Participants were not informed on the neural target, or its association to specific mental processes. Instead, instruction were to freely use mental strategies, allowing individual adoption of most effective strategies. The interface used during most sessions was an animated audio-visual scenario developed and validated previously[36]. During the ’Rest’ condition, a virtual hospital waiting room became more agitated, as the number of virtual characters standing in front of a receptionist increased, along with the loudness of their voices. During the ’Regulate’ condition, Amyg-EFP power was calculated every 3 seconds, and feedback at time *t* was calculated according to the following formula:

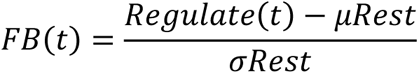

Where μ*Rest* and σ*Rest* are the average and standard deviation of the Amyg-EFP power during the ’Rest’ condition respectively. Successful down-regulation was reflected as lower level of agitation in the room. The PTSD subgroup from Fruchtman et al., [46] had interleaved sessions with an auditory interface (either a musical excerpt or a script of their traumatic experience) during which successful modulation was reflected as lower volume. Key differences between the group protocols are detailed in table 1.

**Table 1.** Amyg-EFP-NF Protocols.

| Sub-Group | Previous Reference | N | No. of Sessions | Interface Type | 'Rest' Length | 'Regulate' Length | No. of Cycles/Session |
| --- | --- | --- | --- | --- | --- | --- | --- |
| HC | Keynan et al., 2019 | 30 | 6 | Audio-visual | 1 min | 1 min | 5 |
| PTSDa | Fruchtman et al., 2021 | 18 | 15 | Auditory/Audio-visual | 1 min | 3 min | 5 |
| PTSDb | Fine et al., 2023 | 20 | 10 | Audio-visual | 1 min | 3 min | 5 |
| PTSDc | -- | 10 | 6 | Audio-visual | 1 min | 3 min | 5 |
| FM | Or-Borichev et al., in prep. | 19 | 10 | Audio-visual | 1 min | 2 min | 8 |

### Online EEG acquisition and processing

EEG data were acquired using the V-Amp EEG amplifier (Brain Products, Munich, Germany) and BrainCap electrode cap mounted with sintered Ag/AgCl ring electrodes. The Pz electrode was positioned according to the standard 10/20 system with a reference electrode placed between Fz and Cz. Raw EEG signal was sampled at 250Hz and broadcasted to the RecView software (Brain Products) enabling custom real time processing through a DLL compiled from Matlab2009b, calculating the Amyg-EFP power every 3 seconds. The Amyg-EFP model was previously developed and validated in our lab to enable the prediction of localized limbic activity from EEG only[42,50].

### EFP capacity calculation

From each session of each participant we calculated an average modulation score using the difference between EFP power during ’Regulate’ and ’Rest’. To achieve a more stable measure for capacity we divided each protocol into three phases – Beginning, Middle and End, averaging together modulation from a number of consecutive sessions. Because of the different training regimens, we used 2 or 3 sessions for each part depending on total length. Amyg-EFP capacity was defined as the modulation during the end of training phase. To account for potential baseline predictors of capacity, the modulation score from the first cycle of the first session was calculated.

### Self-report assessments

Participants were asked to complete translated Hebrew versions of validated self-report questionnaires before and after the training period. Alexithymia was measured using the Toronto Alexithymia Scale (TAS-20), measuring difficulties in expressing and identifying emotions and consisting of 20 items[51]. State anxiety was measured using the STAI questionnaire consisting of 20 items[52]. The Beck Depression Inventory (BDI) was used to measure affective symptoms[53]. A one way ANOVA was performed on each of the psychological measures to examine group differences before training. Change in each measure was calculated as the difference between baseline and post-training score.

### Post training fMRI-NF

To examine the transferability of the capacity acquired during training, each participant performed an assessment of amygdala down regulation in a different context. The fMRI-NF paradigm was of similar design as in the Amyg-EFP procedure, but employed a different interface and was repeated for two consecutive cycles. During the ’Rest’ condition (1 min), an animated figure skated down a rural road at a fixed speed, with a speedometer marking the current speed at all times. During ’Regulate’ (1 min), participants were asked to lower the speed of the skateboard using the same mental strategies they mastered during Amyg-EFP training, with successful down-regulation resulting in a slowing of the environment and marked speed. After each regulate condition, a short summary appeared with the average riding speed (9 sec), followed by a washout fixation period (30 seconds). Of note, the FM group performed a task with the same design during the fMRI-NF, but with an audiovisual interface similar to the one used during training.

A subset of patients from the PTSD and FM groups, performed a brief, similar design fMRI-NF session before starting the Amyg-EFP-NF training course. Target activity during the ’Rest’ condition was used as a measure for baseline reactivity levels.

### fMRI Data Acquisition and Online Feedback Calculation

Structural and functional MRI scans were performed in a 3.0T Siemens MRI system (MAGNETOM Prisma) using a 20-channel head coil. To acquire high-resolution structural images, a T1-weighted three-dimensional (3D) sagittal MPRAGE pulse sequence (TR/TE = 1,860/2.74ms, FA = 8°, voxel size = 1 ×1 × 1 mm, FOV = 256 × 256 mm) was used. Functional whole-brain scans were performed in an interleaved top-to-bottom order, using a T2*-weighted gradient echo planar imaging pulse sequence (HC group: TR/TE = 3,000/35ms, FA = 90°, voxel size = 1.56×1.56×3 mm, FOV = 200×200 mm, 44 slices; PTSD group: TR/TE = 2,500/30ms, FA = 82°, voxel size = 2.3×2.3×3 mm, FOV = 220×220 mm, 42 slices; FM group: TR/TE = 3,000/35ms, FA = 90°, voxel size = 2.3×2.3×3 mm, FOV = 220×220 mm, 46 slices).

During the fMRI-NF session, activity from the right Amygdala was delivered as feedback to the participants. The probing of amygdala-BOLD for NF was based on a 6-mm sphere in Talairach space in the right amygdala (coordinates, 20, -5, -14) in correspondence to the Amygdala BOLD used as a predictor for the AmygEFP model[54]. Momentary beta weights of the predefined ROI (averaged across all voxels of the ROI) were extracted online using Turbo Brainvoyager 3.0 (Brain Innovation). The beta weights were then transferred to MATLAB and feedback was calculated in a same manner as in the Amyg-EFP procedure.

### fMRI data preprocessing and analysis

Imaging data was preprocessed with *fMRIPrep* 21.0.2 [55] which is based on *Nipype 1.6.1* [56]. Anatomical T1 weighted images were corrected for non-uniformity, skull stripped and segmented into CSF, WM and GM tissue. Volume based spatial normalization to standard space (MNI152NLin2009cAsym) was performed through nonlinear registration with ANTs. For the functional images, head motion parameters were estimated with respect to a BOLD reference volume generated using a custom methodology of *fMRIPrep*. Next, volumes were slice-time corrected to the middle of the repetition time, resampled and corrected for head motion. The BOLD reference was co-registered to the T1w reference with boundary-based registration (6 DOF) and preprocessed images were transformed to standard space and smoothed with 5mm FWHM Gaussian kernel.

First level analysis was performed in SPM12 (https://www.fil.ion.ucl.ac.uk/spm/) using four task conditions of interest (’Rest’, ’Regulate’, ’Feedback’, ’Washout’) and additional regressors of no-interest (6 head-motion parameters, averaged BOLD signal from a mask of white matter and csf). Additionally, volumes with a calculated framewise displacement larger than 0.9mm were marked and motion outlier regressors were added to the model (i.e. scrubbing). Statistical maps were estimated for the ’Regulate’>’Rest’ contrast for further analysis.

For each participant an estimate of Amygdala fMRI modulation was extracted using Marsbar toolbox[57]. The right Amygdala region of interest was selected using an anatomical mask based on the Automated Anatomical Labeling atlas[58]. Based on the extracted modulation, high and low capacity participants were defined as those with a negative (n=58) and positive (n=39) beta estimates, respectively.

Next, a second-level analysis was performed using a two samples design, to test the differences between the high and low capacity groups. Peak voxel activations were thresholded at p(FDR=0.005)<0.0007 with clusters larger than 50 voxels.

Finally, a second-level one-sample analysis was performed on the high capacity group, using the modulation score extracted from the amygdala of each participant as a covariate. A whole brain activation map was created from the covariate estimate, showing region co-modulating with the region of interest. Peak voxel activations were thresholded at p(FWE=0.05)<3×10^-7^ with clusters larger than 50 voxels.

Activation clusters from the covariate map were extracted for each participant using the Marsbar toolbox for SPM and used for a regression analysis explaining Amyg-EFP capacity. First, we applied a baseline model using the activation from the right amygdala cluster. Next, we added all activation clusters to a full model. To assess the contribution of each feature in the capacity model, we used the Shapley Additive exPlanations (SHAP)[59] analytical approach. This method is used to explain the output of machine learning models by calculating the contribution of each feature to a particular prediction (sample). Aggregated SHAP scores can reveal the generally more important features across predictions.

For a final assessment of the relationship between the change in alexithymia score and the estimated capacity, individual activations from the significant clusters were weighted according the regression model coefficients, resulting in an estimate of the true capacity.

### Statistical Analysis

All statistical tests were performed using JASP (jasp-stats.org; version 0.9.2). Correlations were performed using Spearman’s correlation coefficient to account for outliers and non-normally distributed data.

## Results

Ninety-seven participants across the three study groups (i.e., HC, FM, PTSD) completed an Amyg-EFP-NF training course (6-15 sessions, see methods), followed by a post-training assessment session that included measurements of their performance with real-time fMRI-NF targeting amygdala down-regulation. Amyg-EFP modulation capacity, which is the focus of this study, was defined as the average signal difference between the ’Regulate’ and ’Rest’ conditions across runs in each training session. Due to diverse training regimens in the different studies and to achieve a stable estimate we calculated for each participant the average capacity across 2-3 sessions at the beginning, middle or end of the training course.

Addressing our first aim of characterizing the establishment of NF capacity, we first tested whether the Amyg-EFP-NF modulation capacity improves over the course of training and across the study groups. A mixed repeated measures ANOVA with training phase as the within subject factor and study group as a between subject factor revealed a significant main effect for training phase (*F*(2) = 3.6, *p* < 0.03), suggesting an overall increase in neuromodulation capacity with additional training (Figure 1B). No study group or study group*training phase effects were found (*group p* > 0.4; *group* ∗ *training p* > 0.6), suggesting no difference in NF regulation across time between a range of disorders, although this cannot be determined definitively from this analysis. As the capacity increases with training, we considered each participants’ end of training score as their gained neuromodulation capacity for further analysis.

Next, we sought to evaluate whether the established Amyg-EFP-NF capacity further extends to the neuromodulation capacity of the amygdala, as depicted by the post training fMRI-NF session. Here, amygdala neuromodulation capacity during fMRI-NF was defined as the contrast in beta estimates between ’Regulate’ and ’Rest’ conditions (Regulate > Rest), which were extracted from the right Amygdala region of interest for each participant. No significant differences were found between the study groups (*F*(2) = 2.25, *p* > 0.1) with a considerable amount of variability noticed within each group. Importantly, linking the two NF-modalities, a significant and positive correlation was found between Amyg-EFP-NF gained neuromodulation capacity (as measured during the end of training period), and the post-training success in the fMRI-NF (Figure 2A, *ρ*(97) = 0.31, *p* = 0.002). This highlights the association between the two measures as indicators of neuromodulation capacity.

**Figure 2.**
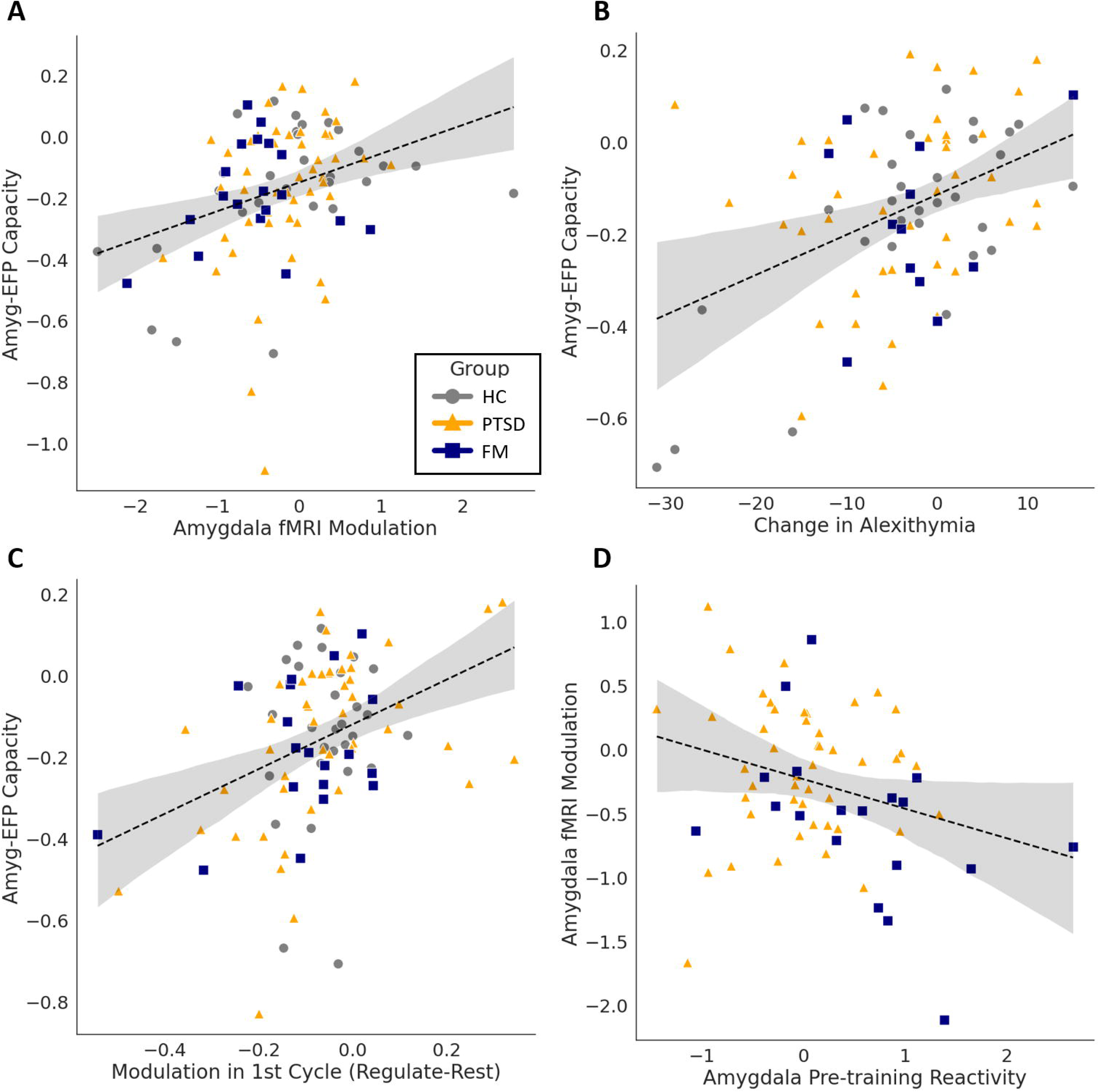
Psychological and Neural correlated of Capacity. Amyg-EFP capacity correlation is shown with (A) Amygdala fMRI modulation during post-training assessment; (B) Change in alexithymia scale score between pre and post-training measurements – negative scores imply improvement in identifying and describing emotions; (C) Amyg-EFP modulation during the first cycle performed. (D) Amygdala fMRI-NF modulation is correlated with pre-training reactivity in a subgroup of patient who performed pre-training NF run. Reactivity is defined as the amygdala activation during the ’Rest’ condition. Across panels, the HC group is shown in gray circles, PTSD in orange triangles and FM in blue squares.

We then turned to test the process-relevance of the established neuromodulation capacity by examining its association with the training-induced changes in Alexithymia scores; an emotion regulation related mental process which is known to involve amygdala activity and was previously demonstrated to be reduced after NF training for down regulating Amyg-EFP[60]. For that, we calculated the difference in the Toronto Alexithymia Scale (TAS) between questionnaires taken before the Amy-EFP training course and after the fMRI session. Negative change scores in this measure suggest an improvement in identifying and describing emotions. As hypothesized, the change in TAS score was associated with Amyg-EFP capacity (Figure 2B, *ρ*(84) = 0.31, *p* < 0.005) suggesting that better regulation capacity is related with greater improvement in alexithymia scores, alluding to process-related neuroplasticity. Amyg-EFP was not associated with changes in any of the other psychological measures (BDI, STAI p’s>0.2)

To complete the characterization of the establishment of regulation capacity, we examined whether there are neurobehavioral tendencies, already measured at baseline, which may predict the extent of gained capacity. For that, we looked for neural factors measured before training, which could predict the achieved capacity. We examined the very first Amyg-EFP modulation cycle, as a measure for the initial capability to modulate the neural target. We found that stronger modulation in the first run was associated with higher Amyg-EFP capacity following training (Figure 2C, *ρ*(94) = 0.358, *p* < 0.001). We then tested the idea that amygdala reactivity, as measured in fMRI can be also a predictor for regulation capacity. The PTSD and FM study group patients had pre-training fMRI scans with a brief NF run. In these datasets, amygdala activation during the ’Rest’ condition was used as a measure for individual baseline reactivity. As hypothesized, we found that reactivity during rest was associated with amygdala fMRI down-modulation in the post-assessment session (Figure 2D, *ρ*(66) = −0.26, *p*=0.03) indicating that higher reactivity level may predict better capacity following training.

Addressing the second aim of this study - to elucidate the neural underpinning of the gained modulation capacity, we divided the entire sample into High and Low capacity sub-groups, according to their success in the fMRI-NF session (considering negative beta estimates as successful modulation and vice versa). Figure 3 shows the whole brain activation maps for the direct comparison between groups (High>Low Capacity, p(FDR=0.005)<0.0007, k>50). The map shows extensive regions beyond the expected right amygdala target, including left amygdala, bilateral hippocampus and parahippocampal gyrus, primary visual areas, bilateral posterior insular cortex, bilateral primary motor cortex, middle cingulate cortex, ventromedial prefrontal cortex as well as basal ganglia (see supplementary table 2). While these results point to large differences between those succeeding and those who do not, the observed differences may be driven not only by variations in processing related to regulation capacity but also by group differences in the perceptual qualities of the interface, whether given mostly positive or mostly negative feedback, or the subjective experience of success.

**Figure 3.**
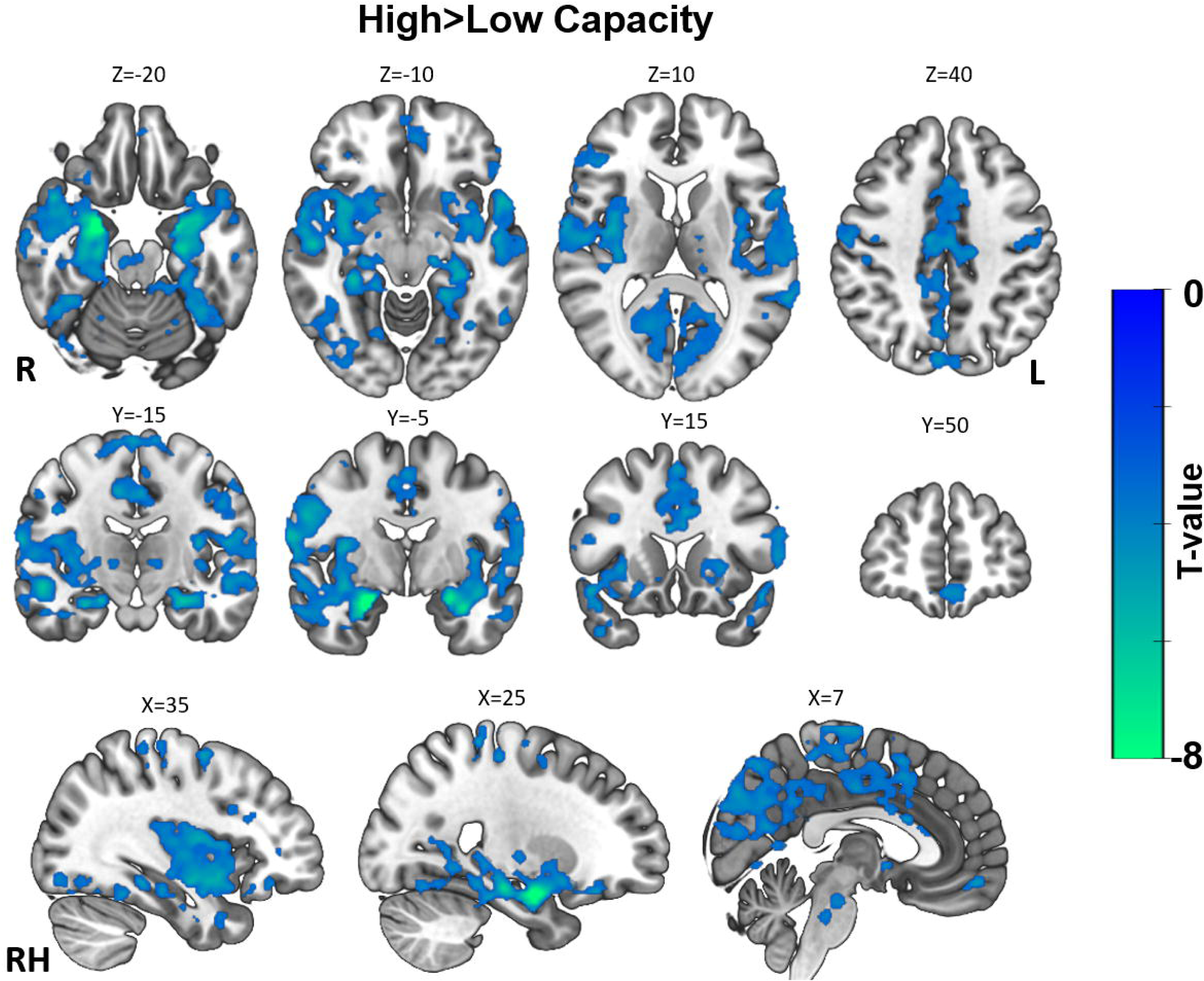
Whole brain differences between High and Low capacity. High and low capacity subjects were defined based on average amygdala modulation, and directly compared using a two-sample second level design. Map shows decreased activity in the High compared to the Low capacity group during the ’Regulate’>’Rest’ contrast, for amygdala down-regulation. Map is thresholded with p(FDR=0.005) and clusters larger than 50 voxels (full table in supplementary materials).

To account for these possible confounds, we further focused on the high-capacity group, and considered the actual capacity from each participant, using a covariate of average amygdala modulation. Figure 4A depicts the whole brain activation map for the ’Regulate’>’Rest’ contrast in the High-capacity sub-group with a second-level capacity covariate derived from the average amygdala modulation. This analysis thus focuses on brain regions co-modulating with the right amygdala, which was the target, during neuromodulation. With a stringent threshold corrected for family-wise error (p(FWE=0.05)<3×10^-7^) a more robust network is introduced, encompassing clusters in the bilateral amygdala, bilateral posterior insula, right parahippocampal gyrus, right cerebellum, right putamen, and right temporal regions (see figure legend and supplementary table 3).

**Figure 4.**
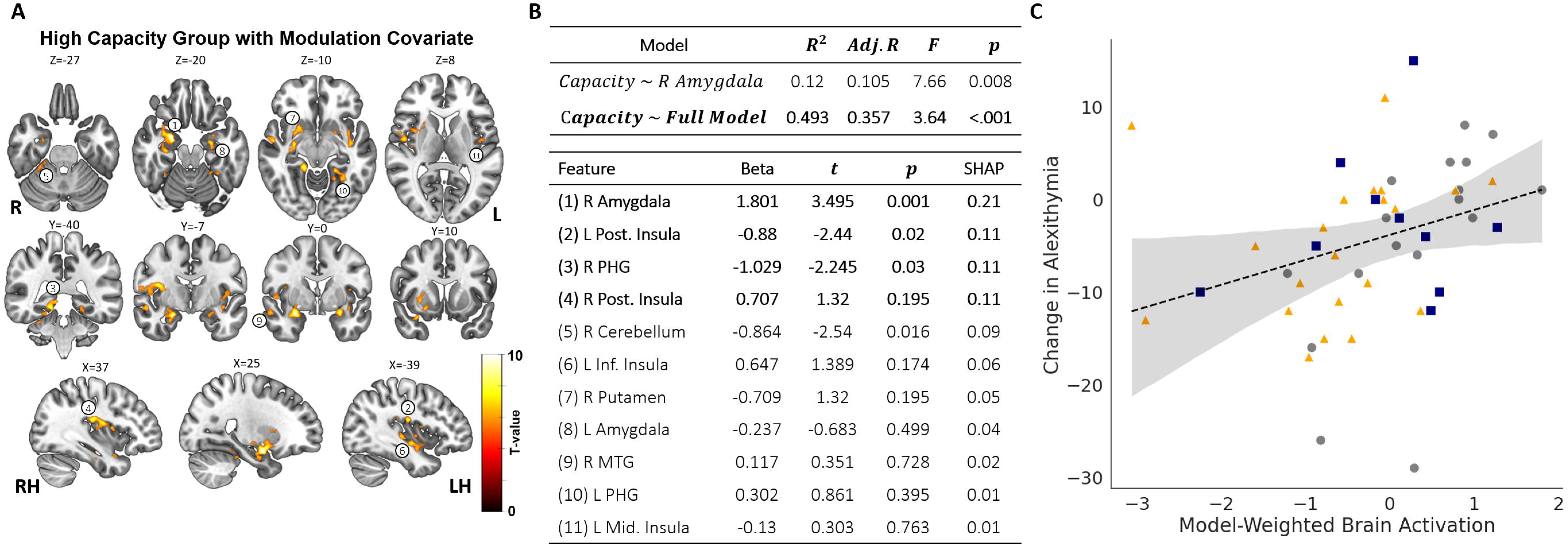
Linear model explaining Amyg-EFP capacity through brain activations. (A) Whole brain activation for the ’Regulate’>’Rest’ contrast in the High capacity group, with a second level modulation covariate based on amygdala modulation. Map is thresholded with p(FWE=0.05) and clusters larger than 50 voxels (full table in supplementary materials). Numbers on the map refer to the full model features. (B) Baseline and full model explaining Amyg-EFP variability, with Amygdala only or all clusters of activation together. Regions in the full model are ordered according to their SHAP values, indicating importance to prediction. (C) Brain activation weighted according to the full model coefficients are associated with the change in alexithymia score (*ρ* = 0.45; *p* = 0.001).

This analysis suggests that the activation in additional regions, extending beyond the amygdala target, may support its established modulation capacity. If these regions’ co-modulation is indeed relevant for capacity, this suggests that their involvement would also explain the variance in the Amyg-EFP capacity. To test this question, we used a linear regression model explaining variability in the Amyg-EFP capacity through neural features extracted from this network. As shown earlier, a basic model including only the right Amygdala target as a feature for Amyg-EFP capacity is significant and explains 10.5% of the variability (adj. R^2^=0.105; F=7.7; p=0.007). Remarkably, when adding all other clusters of activations, the full model increases its explained variability to 35.2% (adj. R^2^=0.35; F=3.58 p<0.001), suggesting that their recruitment is also considerable in explaining the amyg-EFP modulation capacity (Figure 4B). To identify the relative importance of each feature compared with others, importance scores were calculated using the SHAP analytic approach[59]. This method, used for explaining machine-learning models by accounting for the effect of each feature on individual predictions, can rank features in the model by order of average importance. We find that three regions (beyond the right amygdala) with highest SHAP values are the bilateral posterior insula and right parahippocampal gyrus.

Finally, closing the loop from the neuromodulation back to the process, we asked if the highlighted extended network supporting regulation capacity might further explain the association between the Amyg-EFP modulation capacity and reduction in alexithymia scores. Indeed, the weighted activations from the full linear regression model were highly correlated with the reduction in alexithymia score (Figure 4C, *ρ* = 0.45; *p* = 0.001), suggesting that the recruitment of this network may support process engagement.

## Discussion

In the current study, we aimed to thoroughly investigate the neurobehavioral underpinnings of ’neuromodulation capacity’ - the degree of successful volitional self-regulation of a specific neural signal through neurofeedback training. This neuromodulation capacity, which varies widely among individuals, differs from additional aspects of neurofeedback behavior as learning slopes or pre-post changes, and represents an aspect related to the overall aptitude for neural self-regulation. Leveraging a unique dataset from healthy and patient samples, our study combined a relatively extensive Amyg-EFP-NF repeated training period of six to fifteen sessions with post-training Amygdala-fMRI NF sessions. This combination allowed us to both characterize the Amyg-EFP modulation capacity in terms of its association with emotional mental process and neuroanatomical underpinnings (first aim), and unveil the broader neural networks as depicted by fMRI, exceeding the target of modulation (i.e. the amygdala), that are involved in successful Amyg-EFP modulation and process modification (second aim).

### Validation of Amygdala-EFP as a Surrogate for Limbic Neuromodulation Capacity

We first demonstrated, as hypothesized, that Amyg-EFP neuromodulation capacity could be improved through training (Figure 1). Importantly, we also found that such capacity is strongly associated with its neuroanatomical target modulation, as depicted by post-training Amygdala-fMRI-NF; a gold-standard procedure for self-neuromodulation of deeply located limbic regions (Figure 2)[32,61]. Alluding to an associated psychological process, we further showed that the established capacity to downregulate the Amyg-EFP is associated with greater improvement in alexithymia over training. This highlights the process relevance of Amyg-EFP-NF, whereby its acquired regulation capacity, rather than the one observed at start, is associated with improved ability to identify and describe emotions - a process strongly attributed to amygdala function and emotion regulation[62–64]. Demonstrating a reduction in alexithymia following Amyg-EFP-NF is complementing our previous findings among military first responders[23]. It is particularly interesting in light of the alleviated alexithymia scores observed during chronic stress among combat soldiers[65] pointing to a possible stress inoculation effect of learning to downregulate an amygdala-related neural signal. Considering previous research associating alexithymia with post-traumatic stress disorder (PTSD) and combat-related PTSD in particular[66], the current results may further indicate the clinical potential of Amyg-EFP-NF.

These findings collectively support the validity of Amyg-EFP self-neuromodulation as a proxy for limbic modulation across healthy and patient brains. Equally important, these findings highlight the cognitive relevant neuroplastic potential of NF, as reflected by neuromodulation capacity. It shows that self-neuromodulation of limbic activity can be used to induce changes in a relevant mental process of alexithymia, as well as improving the modulation of limbic signal post training; unraveling effects that persist beyond the period of the Amyg-EFP-NF training.

### Pre-existing Neural State and Modulation Success Predict Neuromodulation Capacity

Next, we showed that although neuromodulation capacity is a malleable measure that can improve through repeated training, neurophysiological markers, such as target reactivity at baseline and modulation ability during the very first trial, provide a good predictor of an individual’s full range of neuromodulation capacity. Alongside, and in contrast with prior literature[35], we did not identify any additional predictors, as psychological or symptomatic measures at baseline that predicted neuromodulation capacity as effectively as these neurophysiological markers. This finding alludes to two key issues regarding neuromodulation capacity. First, it points to the relevance of baseline target activation as the foundation for established neuroplasticity (i.e. established capacity). This aligns with a recent meta-analysis, who reported a slight positive correlation between pre-training brain activity in the target regions of fMRI-NF during functional localizer runs and learning success[67]. No other ‘general’ brain-based predictors that are consistent across the diverse studies and targets were found in this study. More broadly, our finding of the importance of the target region reactivity, further corresponds with previous work showing that synaptic activation is important for neural plasticity[68]. This claim has been also put forth with regard to efficacy of other neuromodulation techniques in humans such as Deep Brain Stimulation and Transcranial Magnetic Stimulation. It has been shown that the brain state (i.e. context induced functionality) when the treatment is delivered - is an important factor for stimulation outcome[69]. While activity or state dependency in brain stimulation is a well-acknowledged concept, the mechanisms underlying it have not yet be fully unveiled. The precise relationship between ongoing activity and neuronal excitability is likely complex[70,71], and current studies have yielded opposing effects [for a review see [69]. Our exploratory finding that baseline amygdala activation in the fMRI-NF context was associated with greater fMRI modulation capacity is in line with such claims, though need further investigations. Second, the correlation between neuromodulation ability during the very first attempt and the end-point neuromodulation capacity points to an a priori potential for reaching the maximum neuromodulation capacity. This a-priori ability or disability has been defined as NF literacy[38], though its trait-like nature has been questioned by others[72,73]. Nevertheless, altogether, the findings pointing to the role of pre-existing neural state and trait suggest that personalized preparation for NF might contribute the overall achieved neuromodulation capacity.

### The Neural Basis of Neuromodulation Capacity Extends Beyond the Amygdala Target

The second aim of this study was to investigate the neural underpinning supporting neuromodulation capacity for amygdala down regulation, and to investigate which areas beyond the target (i.e. amygdala) may contribute to the established modulation capacity. For that purpose, we analyzed the post-training amygdala fMRI-NF data. We first characterized the whole brain fMRI patterns that distinguish between individuals with high and low capacity in NF neuromodulation of the amygdala. Next, we searched for regions among those with high capacity, where their co-activation with the amygdala during neuromodulation could best explain the established EFP modulation capacity. Previous accounts for the neural correlates of successful neuromodulation found evidence for a role of additional regions in the prefrontal cortex[74,75] and basal ganglia[76–78], but these were typically limited to within session contrasts, highlighting the reinforcement learning aspects of the task and to a lesser extent, the acquired skill or the established capacity range. Furthermore, these accounts may be further confounded with processes related to the processing of the feedback or success per se[76,79].

The marked categorical differences in brain modulation patterns between high- and low fMRI-capacity groups (Figure 4), previously demonstrated in a single fMRI session with a different sample of healthy participants [16], are replicated here in diverse group of participants (patients and healthy) who underwent extensive Amyg-EFP-NF training prior to the single fMRI-NF session. The results uncovered broad network of regions including the posterior insula, ventromedial prefrontal cortex and hippocampus/parahippocampus. However, the meaning of this broad difference in network recruitment should be interpreted with caution as it could be attributed to general processes related to the NF task, as observed in previous studies[41], or to psychological aspects such as attention allocation and motivation[33]. Furthermore, it could be induced by changes in feedback levels experienced by trainees in the high and low capacity groups, meaning observing different scenarios in the feedback interface. In our dataset, this might be particularly worrisome since the interface used in some of the cases was a complex audio-visual scenario, which became quiet and less densely occupied with avatars during successful modulation[36]. To account for these optional explanation we applied an additional analysis focusing on the high-capacity group in fMRI-NF and used second-level covariate analysis. This analysis identified a more restricted network of regions where neuromodulation, tied to the amygdala, also corresponded with the gained neuromodulation capacity of the Amyg-EFP signal (Figure 5). Using an analytical approach to generate feature importance, we further found that the strongest contributing regions to the prediction were the bilateral posterior insula and the right parahippocampal gyrus. These regions are generally found to be functionally connected to the amygdala at rest [80–82] but, intriguingly, also play an important role for shared mental processing related to emotion regulation such as alexithymia [60,83–85]. Of particular interest is the co-modulation of the amygdala with posterior insula, as both regions are recognized for their roles in processing incoming interoceptive signals related to internal bodily states, such as cardiac and respiratory functions. Notably, an increase in the neural amplification of interoceptive signals, possibly through heightened neural gain of afferent input to these regions, has been linked to an intensified stress response[86]. This finding suggests a potential involvement of interoceptive regulation processes in the success of down regulating the amygdala through NF. However, it is important to acknowledge that such interpretations primarily rely on reverse inference, which infers the presence of specific cognitive processes from observed brain activity patterns. Therefore, we encourage further investigations to build upon these exploratory findings in a hypothesis-driven manner.

### A Network Based-Perspective of Neurofeedback

As previously mentioned, investigating the neural mechanism of neuromodulation capacity with respect to a mental process holds promise for uncovering principles of such cognitive processes. In neurofeedback, when brain activity is manipulated in a neuroanatomically precise manner, it offers insights into the functioning of this brain region and its associated network. Unlike traditional lesion studies, neurofeedback does not rely on external stimulation hardware, allowing trainees to learn to control their brain activity. This approach aids in understanding the broader dynamics of brain function and facilitates the development of mechanistic theories[87]. Similar to recent findings[88], our finding shows that neurofeedback gains significance in the context of connectomics, where manipulating one brain area can impact or recruit a set of distant regions due to the brain’s complex network structure.

The evolving perspective of the brain as a network, supported by empirical evidence spanning various spatial and temporal scales, underscores the potential for novel insights from unveiling the beyond the target of modulation (e.g. the amygdala). We believe that establishing a network framework for neurofeedback capacity is essential for refining feedback paradigms aimed at identifying neurophysiological drivers of cognition, and for treating neurological and psychiatric disorders while minimizing adverse effects[89]. This suggestion aligns with a recent network perspective of neurofeedback that has been proposed within the framework of Network Control Theory (NCT) [90]. In the context of neuroscience, NCT, which originates from control theory in engineering, is based on the principle that altering the activity of one node would trigger repercussions across the entire network. Additionally, it posits that certain nodes exert more influence over the network due to their strategic positions within the overall structure. Accordingly, nodes with significant influence, termed control points, can exert varying impacts on the network. NCT suggests that by pinpointing the appropriate set of control points and determining the optimal method to adjust their activity, it becomes feasible to guide the brain from any starting state to a desired target state by applying specific energy patterns to these crucial nodes.

In light of this framework, our findings suggest that repeated sessions of Amyg-EFP down regulation can potentially train individuals to alter an entire network dynamics, and to possibly manipulate various aspects of brain network activity, such as flexibility or participation in different functional modules. This opens possibilities for investigating how altering certain nodes, with greater influence may exert impact on the network and associated process, such as emotion regulation, which could greatly enhance our understanding of neurological and psychiatric disorders (see an proof of concept of this idea in [91]). Furthermore, adapting feedback signals based not only on the target but also on network statistics could revolutionize intervention strategies for patients with disrupted functional connectivity patterns.

### Strengths and limitations

While our study benefits from using a large dataset of Amygdala down-regulation training performed at the same site with multiple training sessions, some limitations should be noted when considering these findings. The use of two distinct patient groups along with healthy controls may hinder our ability to observe symptom related changes or correlations. We have limited the observation to psychological traits measured with questionnaires in all participants, but cannot rule out the contribution of other factors to neuromodulation capacity such as mood, motivation and attention allocation as mentioned before[33]. The different training regimens (e.g. number of sessions, type of feedback, etc.) may also add variability to our sample, resulting in different capacity levels in the patients who underwent longer training (Figure 1B). The added variability of the different feedback interface schemes (i.e. noisy audio-visual scenario or quiet running figure) may also add some variability to some of the analysis like the baseline predictions and group difference. Future studied should control or modulate these factors to determine whether they are associated with specific effects. Nevertheless, this variability in population, training regime and feedback interface also suggests that the highlighted findings may be more generalizable, surpassing a specific context or state. Our results point to a network of regions harnessed among individuals with high capacity as related to successful neuromodulation, however these are limited to the neural target of amygdala during down regulation. Future studies or meta-analyses should explore the possibility that this network is also recruited during other NF training targets and while interacting with different interfaces.

### Conclusions

This study presents the idea of *neuromodulation capacity* as an NF outcome separated from learning. Our findings suggest that in the case of down regulating the Amyg-EFP, such capacity is associated with improved alexithymia, reflecting better emotion regulation; a critical domain in mental health. The relevance of pre-training target activation and the recruitment of beyond the target network for successful regional modulation points to a possible mechanism of NF induced neuroplasticity. These findings support the idea that NF outcome could be further improved by accounting for a broader brain state before and during the training.

## Acknowledgements

We would like to think the Sagol Famliy fund for their support in this project. This work received funding from the European Union’s Horizon 2020 Framework Programme for Research and Innovation under the Specific Grant Agreement No. 945539 (Human Brain Project SGA3), by the ISRAEL SCIENCE FOUNDATION (grant No. 2923/20), within the Israel Precision Medicine Partnership program.

## Declaration of Competing Interests

Talma Hendler is the Chief Medical Scientist and Chair of advisory board in GrayMatters Health Co Haifa Israel, and has stock options. This company sells the platform of “Prism for PTSD”, neurofeedback driven by amygdala-related biomarker, which is also used as the NF probe in this study. Other authors declare no conflict of interest.

