## Supplemental tables for "Amygdala Self-Neuromodulation Capacity as a Window for Process-Related Network Recruitment"

Supplementary Table 1

**Group demographics and baseline measures**

| Sub-Group | N | Age  [m±sd] | Gender (M/F) | TAS-20 [m±sd] | BDI  [m±sd] | STAI  [m±sd] |
| --- | --- | --- | --- | --- | --- | --- |
| HC | 30 | 19.51±1.5 | 30/0 | 42.3±10.1 | 4.2±3.8 | 31.7±11.1 |
| PTSD | 48 | 36.2±11.3 | 16/32 | 61.1±11.9 | 37.4±13.2 | 58.1±11.7 |
| FM | 19 | 35.2±9.2 | 2/17 | 53.3±13.7 | 15.9±8.6 | 48.7±7.7 |

Supplementary Table 2

**Whole brain analysis of the difference between high and low capacity groups (p(FDR<0.005)<0.0007, k>50)**

| Region | L/R | Peak MNI Coords | | | | k | T | p(FDR-corr) |
| --- | --- | --- | --- | --- | --- | --- | --- | --- |
|  |  | X | Y | | Z |  |  |  |
| Amygdala | R | 21.5 | | -4.5 | -18.5 | 25871 | 11.51 | <0.001 |
| Orbitofrontal Gyrus | L | -32.5 | | 39.5 | -14.5 | 223 | 5.45 | 6x10^-5^ |
| Middle Frontal Gyrus | R | 39.5 | | 21.5 | 21.5 | 162 | 5.17 | 0.0001 |
| Frontal Eye Fields (BA8) | R | 35.5 | | -0.5 | 57.5 | 139 | 5.15 | 0.0001 |
| Ventromedial Prefrontal | R | 3.5 | | 53.5 | -14.5 | 229 | 4.92 | 0.0002 |
| Posterior Superior Temporal Gyrus | L | -54.5 | | -56.5 | 15.5 | 110 | 4.84 | 0.0002 |
| Precentral Gyrus | L | -60.5 | | -30.5 | 37.5 | 51 | 4.80 | 0.0002 |
| Cerebellum | L | -20.5 | | -86.5 | -22.5 | 86 | 4.65 | 0.0003 |
| Ventral Anterior Cingulate | L | -0.5 | | 33.5 | -10.5 | 62 | 4.63 | 0.0003 |
| Inferior Temporal Gyrus | R | 53.5 | | -44.5 | -6.5 | 95 | 4.58 | 0.0004 |
| Superior Frontal Gyrus | R | 53.5 | | 11.5 | 31.5 | 55 | 4.53 | 0.0004 |
| Ventral Tegmental Area | R | 3.5 | | -12.5 | -10.5 | 58 | 4.44 | 0.0005 |
| Midbrain | L | -0.5 | | -20.5 | -24.5 | 133 | 4.40 | 0.0005 |
| Superior Frontal Gyrus | R | -18.5 | | 9.5 | 55.5 | 65 | 4.29 | 0.0007 |
| Precentral Gyrus | L | -50.5 | | -32.5 | 59.5 | 65 | 4.21 | 0.0008 |
| Superior Frontal Gyrus | L | -30.5 | | 29.5 | 43.5 | 73 | 4.18 | 0.0008 |
| Cerebellum | L | -40.5 | | -78.5 | -22.5 | 51 | 4.03 | 0.001 |
| Medial Occipital | R | 15.5 | | -98.5 | 1.5 | 62 | 3.96 | 0.001 |
| Cerebellum | R | 15.5 | | -84.5 | -18.5 | 81 | 3.95 | 0.001 |
| Inferior Parietal Lobule | L | -28.5 | | -68.5 | 35.5 | 78 | 3.84 | 0.001 |

Supplementary Table 3

**Whole brain analysis of the High Capacity Group with Amygdala Modulation as second-level covariate (p(FEW=0.05)<3x10-7, k>50)**

| Region | L/R | Peak MNI Coords | | | | k | T | p(FDR-corr) |
| --- | --- | --- | --- | --- | --- | --- | --- | --- |
|  |  | X | Y | | Z |  |  |  |
| Amygdala | R | 21.5 | | -4.5 | -18.5 | 436 | 15.04 | <0.0001 |
| Posterior Insula | L | -40.5 | | -14.5 | 19.5 | 111 | 8.96 | 1.4x10^-7^ |
| Parahippocampal Gyrus | R | 17.5 | | -38.5 | -8.5 | 119 | 8.88 | 2.04x10^-7^ |
| Posterior Insula | R | 39.5 | | -4.5 | 15.5 | 370 | 8.85 | 2.3x10^-7^ |
| Parahippocampal Gyrus | L | -24.5 | | -52.5 | -2.5 | 242 | 8.70 | 4.04x10^-7^ |
| Amygdala | L | -22.5 | | -10.5 | -22.5 | 129 | 8.38 | 1.35x10^-6^ |
| Inferior Insula | L | -40.5 | | -8.5 | -8.5 | 139 | 7.94 | 7.57x10^-6^ |
| Cerebellum | R | 29.5 | | -46.5 | -22.5 | 54 | 7.46 | 4.87x10^-5^ |
| Putamen | R | 29.5 | | 9.5 | -0.5 | 81 | 7.41 | 5.86x10^-5^ |
| Middle Temporal Gyrus | R | 57.5 | | -0.5 | -16.5 | 77 | 7.24 | 0.00011 |
| Superior Temporal Gyrus | R | 65.5 | | -18.5 | 15.5 | 91 | 7.19 | 0.00013 |
| Middle Insula | L | -42.5 | | -8.5 | 3.5 | 52 | 6.86 | 0.00048 |
